# The Arabidopsis R-SNARE VAMP714 is essential for polarization of PIN proteins in the establishment and maintenance of auxin gradients

**DOI:** 10.1101/821272

**Authors:** Xiaoyan Gu, Kumari Fonseka, Stuart A. Casson, Andrei Smertenko, Guangqin Guo, Jennifer F. Topping, Patrick J. Hussey, Keith Lindsey

**Affiliations:** Department of Biosciences, Durham University, South Road, Durham DH1 3LE, UK; Ministry of Education Key Laboratory of Cell Activities and Stress Adaptations, School of Life Sciences, Lanzhou University, Lanzhou 730000, China; Department of Crop Science, Faculty of Agriculture, University of Ruhuna, Mapalana, Kamburupitiya, Sri Lanka; Department of Molecular Biology and Biotechnology, University of Sheffield, Firth Court, Western Bank, Sheffield S10 2TN, UK; Institute of Biological Chemistry, Washington State University, Pullman WA 99164, USA

**Keywords:** *Arabidopsis thaliana*, auxin transport, PIN proteins, R-SNARE, VAMP714

## Abstract

- The plant hormone auxin and its directional intercellular transport plays a major role in diverse aspects of plant growth and development. The establishment of auxin gradients in plants requires asymmetric distribution of members of the auxin efflux carrier PIN-FORMED (PIN) protein family to the plasma membrane. An endocytic pathway regulates the recycling of PIN proteins between the plasma membrane and endosomes, providing a mechanism for dynamic localization.
- N-ethylmaleimide-sensitive factor adaptor protein receptors (SNAP receptors, SNAREs) mediate fusion between vesicles and target membranes and are classed as Q- or R-SNAREs based on their sequence. We analysed gain- and loss-of-function mutants, dominant negative transgenics and protein localization of the Arabidopsis R-SNARE VAMP714 to understand its function.
- We demonstrate that VAMP714 is essential for the insertion of PINs into the plasmamembrane, for polar auxin transport, and for root gravitropism and morphogenesis. *VAMP714* gene expression is upregulated by auxin, and the VAMP714 protein co-localizes with ER and Golgi vesicles and with PIN proteins at the plasma membrane.
- It is proposed that VAMP714 mediates the delivery of PIN-carrying vesicles to the plasma membrane, and that this forms part of a positive regulatory loop in which auxin activates a VAMP714-dependent PIN/auxin transport system to control development.

## Introduction

The polarity of eukaryotic cells is associated with diverse aspects of cell differentiation and development, and one feature of this is the polar distribution of membrane proteins, such as to promote directional signalling or transport of molecules or ions. In plants, local biosynthesis and the regulated polar transport of auxin contribute to the generation of auxin gradients within tissues, necessary for spatially regulated gene expression and development (Reinhardt *et al*., 2000; Petrasek & Friml, 2009; Vanneste & Friml, 2009). Members of the PIN-FORMED (PIN) family of auxin efflux carriers accumulate in the plasma membrane on specific sides of the cell and determine the direction of auxin flow through tissues (Wiśniewska *et al*., 2006; Vieten *et al*., 2007).

Rapid changes in cell polarity involve clathrin-mediated endocytosis of PINs, dependent on both ARF-GEF (guanine-nucleotide exchange factors for ADP-ribosylation factor GTPases)- and Rab5 GTPase-dependent recycling (Steinmann *et al*., 1999; Geldner *et al*., 2001; Kleine-Vehn *et al*., 2008; Kitakura *et al*., 2011). Auxin itself inhibits this recycling, resulting in an accumulation of PIN proteins at the plasmamembrane, so promoting its own efflux (Paciorek *et al*., 2005). While the endocytic model accounts for the dynamic mobilization of PINs to different surfaces of the cell, it does not explain mechanistically how PIN proteins are delivered to the plasma membrane following their translation in the endoplasmic reticulum (ER).

Eukaryotes have evolved N-ethylmaleimide-sensitive factor adaptor protein receptors (SNAP receptors, SNAREs) as mediators of fusion between vesicular and target membranes. SNAREs can be grouped as Q- and R-SNAREs based on the occurrence of either a conserved glutamine or arginine residue in the centre of the SNARE domain (Fasshauer *et al*., 1998). In Arabidopsis, Vesicle-Associated Membrane Protein7- (VAMP7)-like R-SNAREs fall into two gene families - four VAMP71 group proteins are involved in endosomal trafficking (Uemura *et al*., 2004; Hong 2005) and eight VAMP72 group proteins are involved in secretion (Sanderfoot, 2007; Zhang *et al*., 2015). VAMPs have roles in abiotic stress tolerance (VAMP711, VAMP712; Leshem *et al*., 2010; Xue *et al*., 2018), in gravitropic responses (Yano *et al*., 2003), in cell plate formation (VAMP721, VAMP722; Zhang *et al*., 2011; Karnik *et al*., 2013; EI-Kasmi *et al*., 2013; Yun *et al*., 2013; Zhang *et al*., 2017; Uemura *et al*., 2019), in cytokinesis (Collins *et al*., 2003; Karnik *et al*., 2013), in defence responses (Kwon *et al*., 2008; Zhang *et al*. 2011, 2017), and in the transport of phytohormones (Dacks *et al*., 2002; Enami *et al*., 2009).

We identified a gain-of-function mutant of *VAMP714* following an activation tagging screen in Arabidopsis (Casson & Lindsey, 2006). VAMP714 is structurally related to VAMPs 711, 712 and 713, and previous data indicate that, while GFP fusions with VAMP711, 712 and 713 localize to the vacuole in Arabidopsis suspension culture protoplasts, GFP-VAMP714 co-localizes with the Golgi marker VENUS-SYP31 (Uemura *et al*., 2004), but its function is unknown. In this paper, we use a combination of genetics, trangenics and cell biological approaches to investigate the function of VAMP714.

## Materials and methods

### Plant materials

Wildtype *Arabidopsis thaliana* plants (ecotype Col-0) and activation tagging populations (Casson & Lindsey, 2006) and growth conditions (Casson *et al*., 2009) have been described previously. We identified two loss-of-function mutants of *VAMP714* from the SALK and GABI-Kat collections of T-DNA insertion mutants (SALK_005917 and GABI_844B05; www.signal.salk.edu), obtained from the Nottingham Arabidopsis Stock Centre (Nottingham University, Sutton Bonington, UK). RT-PCR analysis showed that neither mutant expresses the *VAMP714* gene to detectable levels. PCR was used to identify homozygous insertion mutants among the GABI_844B05 F1 plants, using oligonucleotide primers to amplify the *VAMP714* gene from wildtype but not from insertion lines: 5’-CTGTTGTAGCGAGAGGTACCG-3’ and 5’-AAGCATGTCAACAAGACCCTG-3’. To confirm T-DNA insertion sites, a *VAMP714* primer (5’-AAGCATGTCAACAAGACCCTG-3’) and a T-DNA left border primer (5’-ATATTGACCATCATACTCATTGC-3’) were used to amplify the T-DNA flanking sequence.

Genetic crosses between Arabidopsis plants were made under a Zeiss STEMI SV8 dissecting stereomicroscope (Carl Zeiss Ltd., Welwyn Garden City, Herts, UK) as described (Souter *et al*., 2002). Arabidopsis seeds transgenic for the marker QC25 and *DR5::GUS* were kindly provided by Prof. Ben Scheres (Wageningen University).

For hormone/inhibitor treatments of seedlings grown *in vitro,proVAMP714::GUS* seedlings were germinated aseptically on growth medium and at 7 dpg were transferred to growth medium containing auxin (indole-3-acetic acid, IAA) and, for comparison, cytokinin (benzylaminopurine, BAP), the ethylene precursor ACC or the polar auxin transport inhibitor 2, 3, 5-triiodobenzoic acid (TIBA) for a further 5 days before analysis. For drug treatments, five-day-old seedlings were incubated in half-strength MS liquid medium supplemented with 50 μM brefeldin A (50 mM stock in DMSO; Sigma-Aldrich), and 20 μM latrunculin B (20 mM stock in DMSO; Sigma-Aldrich). DMSO in the same final concentration (0.1%) was added to negative controls. Each treatment for confocal imaging was repeated at least three times with similar results.

### Gravitropism assays

Mutant and wildtype seedlings were grown on standard agar plates for 4 days and turned to a 90° angle to measure the angle of bending towards gravity. The angle towards the gravity was measured after 8, 12 and 24 h. The curvature of 20 seedlings for each genotype was determined.

### Gene constructions, plant transformation and transient gene expression

To create dominant negative mutant proteins, we expressed a non-functional fragment of the VAMP714 protein expected to bind to the Qa, Qb and Qc complex of SNARE and inhibit the binding of the native protein (Tyrrell *et al*., 2007). For constructing the dominant negative gene construct, the longin and SNARE domains of the *VAMP714* gene sequence were amplified without the transmembrane domain, using the oligonucleotide primers 5’-GGGGACAAGTTTGTACAAAAAAGCAGGCTTCGTTGTAGCGAGAGGTACCGTG-3’, and 5’-GGGGACCACTTTGTACAAGAAAGCTGGGTCCTATTAGCATTTTTCATCCAAAG-3’. The amplified sequence was cloned directly into the pCR^®^2.1-TOPO vector (Invitrogen, Paisley, UK) and then as an *EcoRV* fragment into the pDNOR207 Gateway entry vector and then pMDC43 destination vector (Invitrogen, Paisley, UK), under the transcriptional control of the CaMV35S gene promoter. qRT-PCR showed that the relative abundance of the truncated transcript of *VAMP714* was higher in dominant negative transgenics than is the native transcript in Col-0 wild type plants (Fig. S1). T4 transgenics were produced by selfing, and at least 10 independent lines were analysed phenotypically.

To amplify the *VAMP714* promoter, the following oligonucleotide primer pairs were used: 5’-GTCGAGCAGAGATCCTAGTTAGTGAGTCC-3’ and 5’-GTCGAGGTGATTCGATGACAGAGAGTGGAG-3’; the promoter PCR product was cloned into pCR^®^2.1-TOPO and then as a *Sal1* fragment into promoterless GUS reporter binary vector pΔGUSCIRCE for *proVAMP714::GUS*.

For the *pro35S::VAMP714:GFP* fusion protein, the coding region was amplified using primers 5’-TTAATTAACGCGATTGTCTATGCTGTTGTAGCG-3’ and 5’-CAGATTTTAAGATCTGCATGATGG-3’, and the product was cloned into the pBIN-GFP vector (Dr. David Dixon, Durham University). For *proVAMP714::VAMP714:CFP* and *proVAMP714::VAMP714:mCherry*, a ca. 3.5 kb genomic fragment, comprising ca. 2 kb promoter and 1.5 kb coding sequence of the *VAMP714* gene, was amplified using primers 5’-GGGGACAAGTTTGTACAAAAAAGCAGGCTCAGAGATCCTAGTTAGTGAGTCC-3’ and 5’-GGGGACCACTTTGTACAAGAAAGCTGGGTCAGATCTGCATGATGGTAAAGTG-3’. The PCR product was cloned into pCR^®^2.1-TOPO vector and then as an *EcoRV* fragment into the pDNOR207 Gateway entry vector and then pGHGWC destination vector. All constructs were validated by sequencing.

Transgene plasmids were introduced into *Agrobacterium tumefaciens* C58C3 by triparental mating, and plant transformation was performed by the floral dip method (Clough & Bent, 1998). Transformed plants were selected using standard growth medium supplemented with kanamycin (50 μg/ml for *proVAMP714::GUS*), Basta (15 μg/ml for *pro35S::VAMP714:GFP*) or hygromycin (50 μg/ml for *proVAMP714::VAMP714:CFP* and *proVAMP714::VAMP714:mCherry*).

Transient expression of *pro35S::VAMP714::GFP* and *ST–RFP* constructs was carried out in onion epidermal peels following microprojectile bombardment using the Helios Gene Gun system (Bio-Rad Laboratories, Hemel Hempstead, UK). Plates containing bombarded onion sections were covered with aluminium foil and incubated at 22° C overnight, after which the inner layer of the onion tissue was peeled off carefully and mounted on a glass slide with drop of water, covered with a coverslip and viewed under a Leica SP5 Laser Scanning Microscope (Leica Instruments, Heidelberg, Germany).

### Gene expression analysis

Localization of GUS enzyme activity in transgenic plants containing the *proVAMP714::GUS* reporter gene was performed as described (Short *et al*., 2018). Stained samples were fixed in Karnovsky’s fixative (4% paraformaldehyde and 4% (v/v) glutaraldehyde in 0.1 M phosphate buffer), dehydrated in an ethanol series and embedded in LR White resin (Historesin™ Embedding Kit, Leica Instruments, Heidelberg, Germany) prior to sectioning, as described (Topping *et al*., 1997).

For transcript analysis, RNA was extracted from seedlings 7 dpg using the RNeasy Plant RNA Extraction kit (Qiagen Ltd., Surrey, UK). RT-PCR was performed using the OneStep RT-PCR kit (Qiagen) as per the manufacturer’s instructions. Oligonucleotide primer pairs used for amplification of *VAMP714* were: 5’-GTCGAGCAGAGATCCTAGTTAGTGAGTCC-3’ and 5’-GTCGAGGTGATTCGATGACAGAGAGTGGAG-3’ primers. The *ACTIN2* gene was used as a positive control, using primers 5’-GGATCGGTGGTTCCATTCTTGC-3’ and 5’-AGAGTTTGTCACACACAAGTGCA-3’.

For quantitative RT-PCR, the following primers were used: for *VAMP714*, 5’-GAGATTCGATCGGTCATGGT-3’ and 5’-GGTAAAGTGATTCCTCCG-3’; for *VAMP713*, 5’-TTGTGAAAACATATGGCCGA-3’ and 5’-CTAGCAACTCCAAACGCTCC-3’; for *VAMP712*, 5’-AACGTACTGATGGCCTCACC-3’ and 5’-ATGTTCGCGGTTTTATCGAC-3’; for *VAMP711*, 5’-GGTGGAGAAACTGCAAGCTC-3’ and 5’-ACACACTTCGCAAAGCAATG-3’; for *IAA1*, 5’-GGAAGTCACCAATGGGCTTA-3’ and 5’-GAGATATGGAGCTCCGTCCA-3’; and for *IAA2*, 5’-CACCAGTGAGATCTTCCCGT-3’ and 5’-AGTCTAGAGCAGGAGCGTCG-3’.

### Auxin transport assays

Basipetal shoot auxin transport assays were carried out as described (Chilley *et al*., 2006). 2.5 cm of inflorescence stem segments lacking branches were excised and the apical (upper) end placed in 20 μl MS salts medium in Eppendorf tubes. This pre-treatment prevents air bubbles entering the auxin transport system. Stem segments were then transferred to fresh tubes containing medium supplemented with 0.08 μCi/ml ^3^H-IAA (approx 3.5 μM IAA), again with the apical ends in the liquid medium. Samples were incubated for 18 hours before the basal 5 mm of the sample was removed and placed in 4 ml of scintillation fluid, and incubated for 48 hours before scintillation. Non-inverted samples (in which the basal ends were placed in the medium) were included to control for non-specific transport. Samples incubated in non-radioactive medium were used to detect baseline activity or radioactive contamination.

Acropetal root auxin transport assays were carried out on 2 dpg Arabidopsis seedlings. Agar blocks (1 % w/v, 2-3 mm wide) were prepared containing 500 nM ^3^H-IAA (specific activity is 5.75 μCi/ml; GE Amersham UK) plus 10 μM IAA in 1% v/v DMSO. The ^3^H-IAA blocks were placed onto the top of roots just below the root-shoot junction. For each root analysed, the distance between the application site and the root tip was constant; the plants were inverted and left for 1 hour per cm. The distal 5 mm at the root tip was removed and the sample transferred to 4 ml scintillation fluid (EcoScint A, National Diagnostics) and incubated for 48 hours before measuring in the scintillation counter.

All data were expressed as distintegrations per minute (DPM). Results represent the means of five independent assays ± SD.

### Protein localization and confocal microscopy

PIN protein immunolocalization was carried out as described (Short *et al*., 2018). Fluorescence levels were quantified using ImageJ software (National Institutes of Health, http://rsb.info.nih.gov/ij). At least three independent analyses were carried out, and for each, six random samples for each of 10 roots were measured, using identical confocal settings for each analysis. Results are presented as means ± standard deviation. We thank Prof. Klaus Palme (University of Freiburg) for kindly donating PIN antibodies. Confocal imaging used a Leica SP5 Laser Scanning Microscope (Leica, Heidelberg, Germany). Light microscopy used a Zeiss Axioscop microscope (Carl Zeiss Ltd, Welwyn Garden City, UK) with DIC/Nomarski optics or an Olympus SZH10 microscope system (Olympus Microscopes, Southend-on-Sea, UK).

## Results

We used an activation tagging screen (Casson & Lindsey, 2006) to identify Arabidopsis mutants defective in root development, and one was associated with the upregulation of gene At5g22360, encoding the 221 amino acids VAMP714 protein - this gene was then the focus of further studies (Fig. 1a). To confirm a potential role of the *VAMP714* gene in root development, two independent loss-of-function insertional mutants were identified from the SALK and GABI-Kat collections of T-DNA insertion mutants (SALK_005917 and GABI_844B05; www.signal.salk.edu), and a dominant negative mutant was constructed (Fig. S1). PCR-based genotyping was used to confirm the sites of T-DNA insertion in the SALK and GABI-Kat lines. In the SALK mutant the T-DNA was located in the first intron, and in the GABI-Kat mutant the T-DNA was located in the third exon. The dominant negative gene construct was designed to comprise the Longin and SNARE domains but lack the transmembrane domain, so that it would bind to the Qa, Qb and Qc complex of SNARE but inhibit the binding of the native protein (Tyrrell *et al*., 2007; Fig. S1).

**Figure 1.**
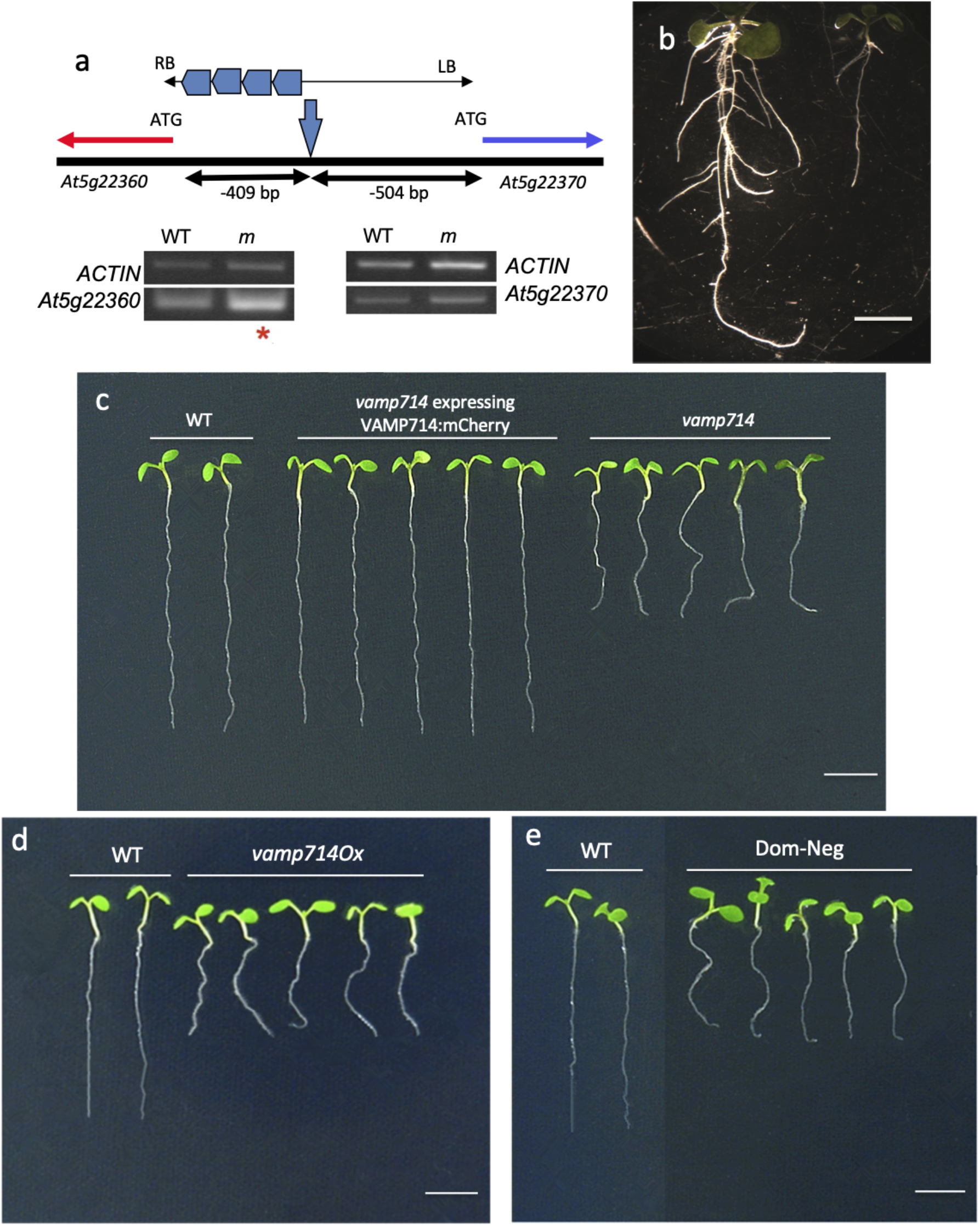
*VAMP714* gene is required for correct seedling development. (a) Diagrammatic representation of the activation tag locus, showing the position of the activation tag T-DNA and the expression analysis of the *At5g22360* and *At5g22370* genes relative to the *ACTIN2* and Col-0 wild type. RB and LB indicate the borders of the T-DNA insertion element, encompassing enhancer regions (the four pentagons) and showing the site of insertion. The distances from the T-DNA insertion site to the transcriptional start codon of each of the adjacent genes are indicated. Red asterisk indicates enhanced expression of *At5g22360* (*VAMP714*) in the mutant (m), compared to in wildtype (WT). (b) Wildtype (left) and activation-tagged *VAMP714* overexpressing (right) seedlings at 14 dpg. Bar = 5 mm. (c) Seedlings (7 dpg) of wildtype (WT), *vamp714* mutant and *vamp714* mutant tranformed with a *proVAM714P::VAMP&14:mCherry* gene fusion, showing functional complementation of the mutant by the fusion gene. Bar = 5 mm. (d) Seedlings (7 dpg) of wildtype (WT) and *pro35S::VAMP714* transgenic overexpressers. Bar = 5 mm. (e) Seedlings (7 dpg) of wildtype (WT) and VAMP714 dominant-negative mutant transgenics. Bar = 5 mm.

Seedlings of both *vamp714* mutants each showed very similar phenotypes, and were smaller than wildtype, with reduced root systems (Figs. 1b,c). The mutant phenotype was functionally complemented by a *proVAMP714::VAMP714:mCherry* transgene (Fig. 1c), showing that the correct gene had been identified, corresponding to the phenotype. By 21 dpg *vamp714* insertional mutants grown in soil developed a shorter primary root than wildtype (1.2 ± 0.2 cm versus 3.5 ± 0.5 cm, n = 20; Fig. 2a), with fewer lateral roots (3.1 ± 1.0 versus 10.0 ± 2.0, n = 20; Fig. 2b) though this represents only a slightly reduced lateral root density (a mean of 2.6 lateral roots per cm at 21 dpg for *vamp714* compared with 2.8 for wildtype). Both transgenic *VAMP714* overexpressers (Fig. 1d) and dominant-negative mutants also showed a reduced root system (Fig. 1e), with a mean primary root length of 1.8 ± 0.2 cm (n = 20) at 21 dpg, and mean lateral root numbers of 3.2 ± 0.9, n = 20. Compared to wildtype, the transgenic overexpressors, *vamp714* loss-of-function mutants and dominant negative mutant plants each showed a dwarfed and excessive leaf and shoot branching phenotype (Fig. 2c-f). As seen in other systems, the phenotypic similarities between plants with loss-of-function and gain-of-function (over-/mis-expressors) of VAMP714 may be due to the disruption of interaction with partner proteins in both mutants and over-/misexpressors (reviewed by Prelich, 2012), and this observation suggests that the stoichiometry of protein complexes in which VAMP714 in involved is required for correct function.

**Figure 2.**
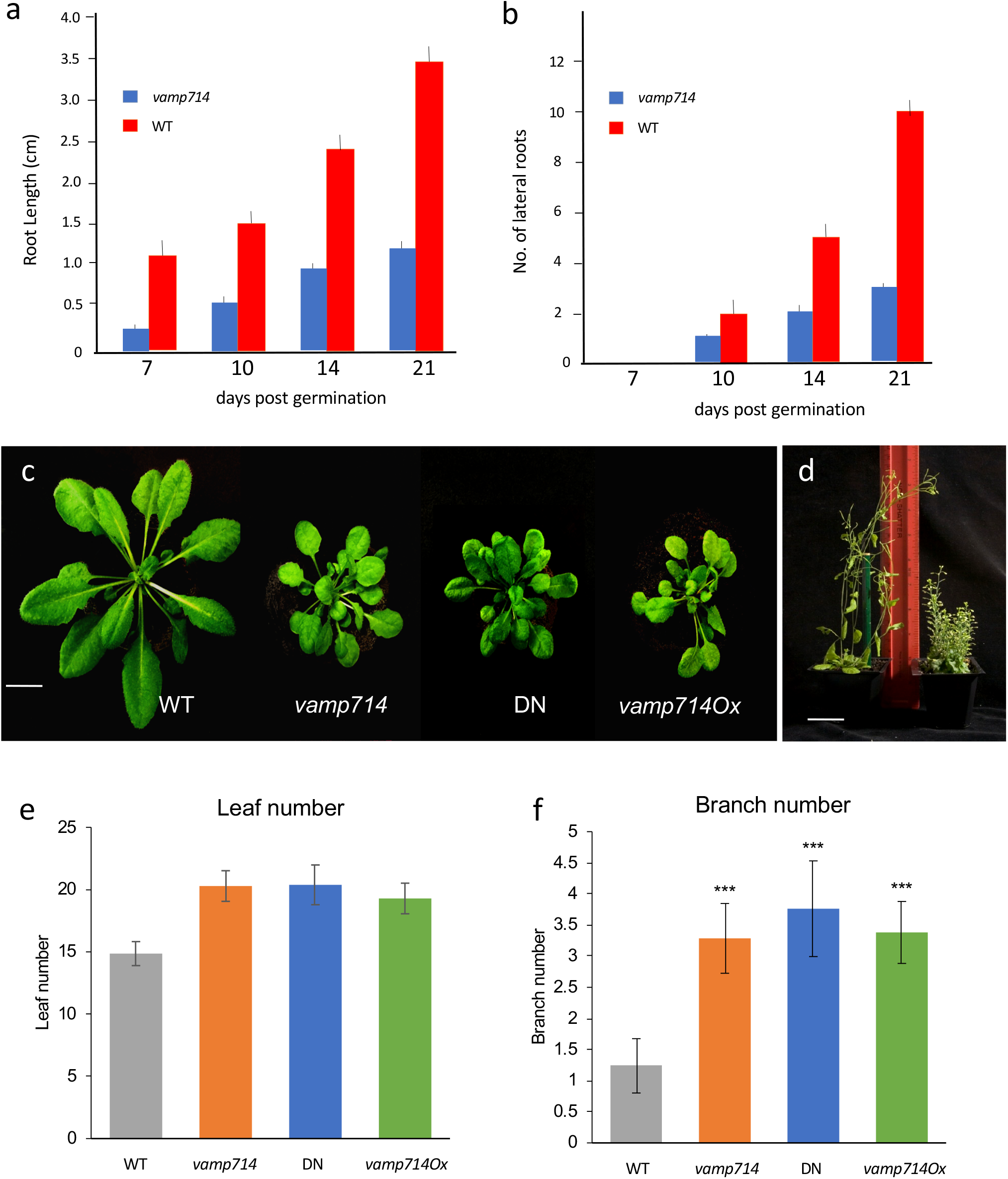
*VAMP714* gene is required for correct root and shoot architecture. (a) Primary root length of wildtype (WT) and *vamp714* loss-of-function mutants grown on vertical agar plates over 21 dpg. Mean of 20 replicates ± standard error of the mean. (b) Lateral root number of wildtype (WT) and *vamp714* loss-of-function mutants grown on vertical agar plates over 21 dpg. Mean of 20 replicates ± standard error of the mean. (c) Shoot phenotypes of wildtype (WT), *vamp714* mutant, VAMP714 dominant-negative mutant (DN) and transgenic *VAMP714* overexpressers (VAMPOx) seedlings at 4 weeks post germination. Bar = 1 cm. (d) Wildtype (L) and transgenic *pro35S:VAMP714* plants (R) plants at 8 weeks post germination. Bar = 3 cm. (e) Rosette leaf number of wildtype (WT), *vamp714* mutant, VAMP714 dominant-negative mutant (DN) and transgenic *VAMP714* overexpressers (VAMPOx) seedlings at 4 weeks post germination. Error bars represent standard deviation of the mean of 3 biological replicates. (f) Shoot branch number of wildtype (WT), *vamp714* mutant, VAMP714 dominant-negative mutant (DN) and transgenic *VAMP714* overexpressers (VAMPOx) seedlings at 8 weeks post germination. Error bars represent standard deviation of the mean of 3 biological replicates. *** indicates significant difference at *P* < 0.005, Student’s *t*-test.

Propidium iodide staining of *vamp714* mutant roots reveals a more disorganized tissue patterning compared with Col-0 (Fig. 3a-d). Lugol staining of *vamp714* mutant roots similarly showed an abnormal patterning of starch grain-containing columella cells, lacking both the discrete columella tier delineation seen in the wildtype and specification of the quiescent centre (QC) - *vamp714* mutants lack an appropriately specified QC and the columella stem cells showed evidence of differentiation (starch accumulation), suggesting a failure of QC activity (Fig. 3e-h). To further investigate QC and stem cell gene expression in these plants, we measured the transcription of the genes *SHORTROOT (SHR*) and *SCARECROW (SCR*) (Sabatini *et al*., 2002) at 7 dpg by qRT-PCR. The transcript levels of both genes were reduced in *vamp714* insertional mutants, dominant negative mutants and overexpressers, consistent with the loss of identity of QC cells and possibly of other stem cells in which these genes are expressed (Fig. 3i).

**Figure 3.**
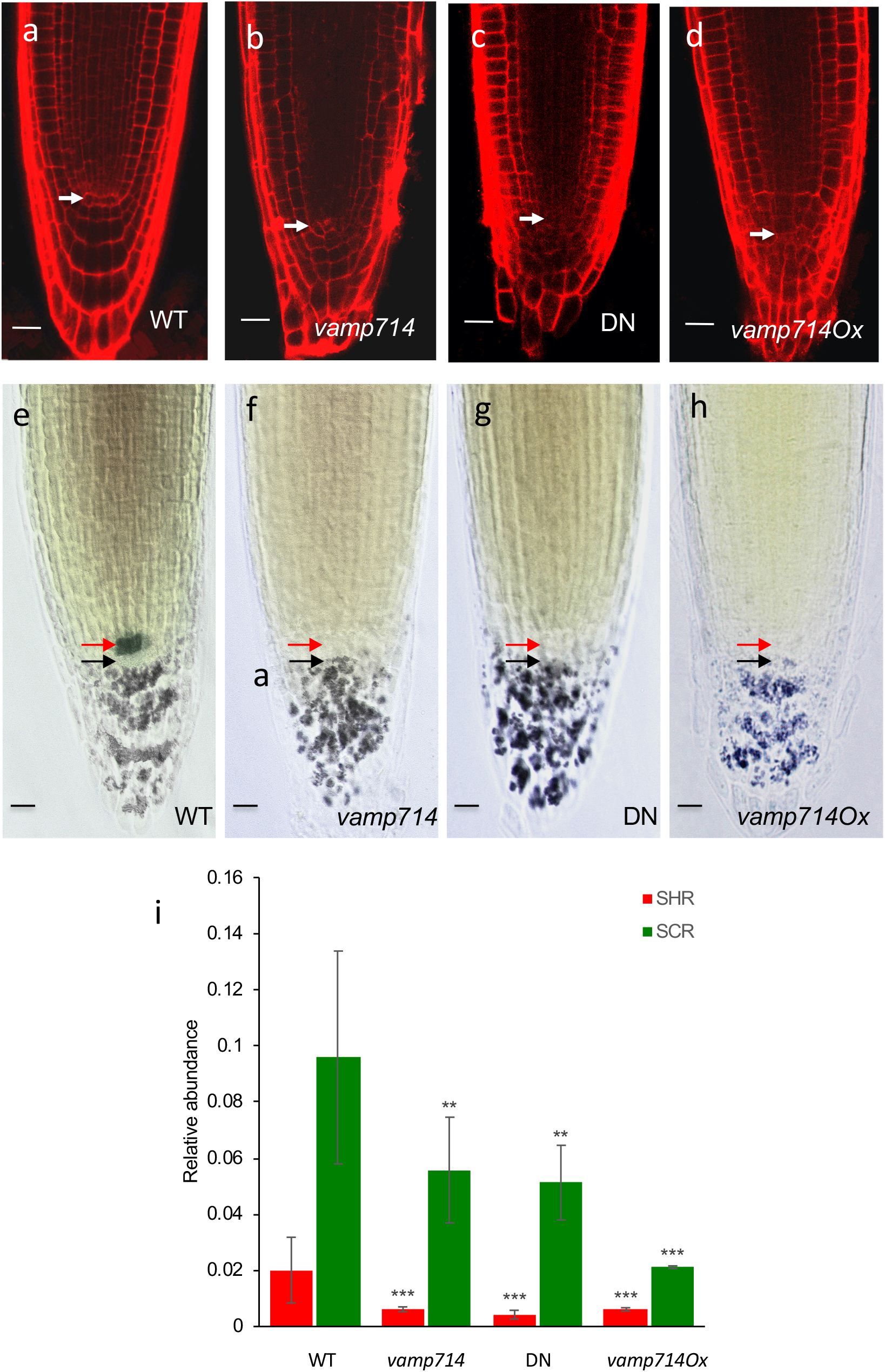
Functional VAMP714 is required for correct root cell patterning, QC maintenance and meristem gene expression. (a-d) Confocal imaging of root tips of (a) wildtype (WT), (b) *vamp714* mutant, (c) VAMP714 dominant negative mutant (DN) and (d) transgenic VAMP714 overexpressing (vamp714Ox) seedlings (7 dpg) stained with propidium iodide. White arrows indicate position of the QC cells. Bars = 10 uM. (e-h) Double labeling of QC and differentiated columella cells visualized by QC25 and amyloplast (lugol) staining, respectively, in wildtype (a) and mutant or overexpressing seedlings (f-H) revealing defective columella stem cells (black arrows) and lack of QC marker expression in *vamp714* mutant (f), VAMP714 dominant negative mutant (g) and *VAMP714* transgenic overexpressing roots (h), showing defects in QC identity (red arrows) and columella patterning. Bars = 10 uM. (i) qRT-PCR analysis of mRNA abundance of the QC identity genes *SHR* and *SCR* in wildtype (WT), *vamp714* mutants, dominant negative mutants (DN) and transgenic *VAMP714* overexpressers (*vamp714Ox*), compared to *ACTIN2* expression. Data represent means of 3 biological replicates ± SD. Significant differences between wildtype and mutant expression at *P* < 0.05 (**) and 0.005 (***) are indicated, Student’s *t*-test.

Consistent with the predicted role as a vesicle-associated membrane protein, a VAMP714:GFP fusion (under the transcriptional control of the *CaMV35S* gene promoter) was constructed for testing *in vivo*, and found to locate to vesicles. Given that a VAMP714:mCherry fusion protein is functional, as demonstrated by genetic complementation (Fig. 1c), we expect a VAMP714:GFP fusion to similarly be functional. Stably transformed Arabidopsis plants expressing *pro35S::VAMP714:GFP*, and transiently transformed onion epidermal peels or tobacco leaf, show GFP signal in discrete vesicles, with additional plasma membrane localization seen in the stable transformants (Fig. 4). The vesicles were identified as Golgi by co-labelling with the Golgi membrane marker ST–RFP (sialyltransferase–red fluorescent protein, Runions *et al*., 2006; Fig. 4a-d) and also some co-localization with the endoplasmic reticulum-targeted red fluorescent protein RFP-HDEL (Lee *et al*., 2013; Fig. 4e-g) and at the plasmamenbrane (Fig. 4h). This is consistent with computational and previous experimental predictions of subcellular location in Arabidopsis (Uemura *et al*., 2004; Fig. 4i). We showed by video confocal microscopy that the vesicles are mobile (Fig. 4h and Video S1).

**Figure 4.**
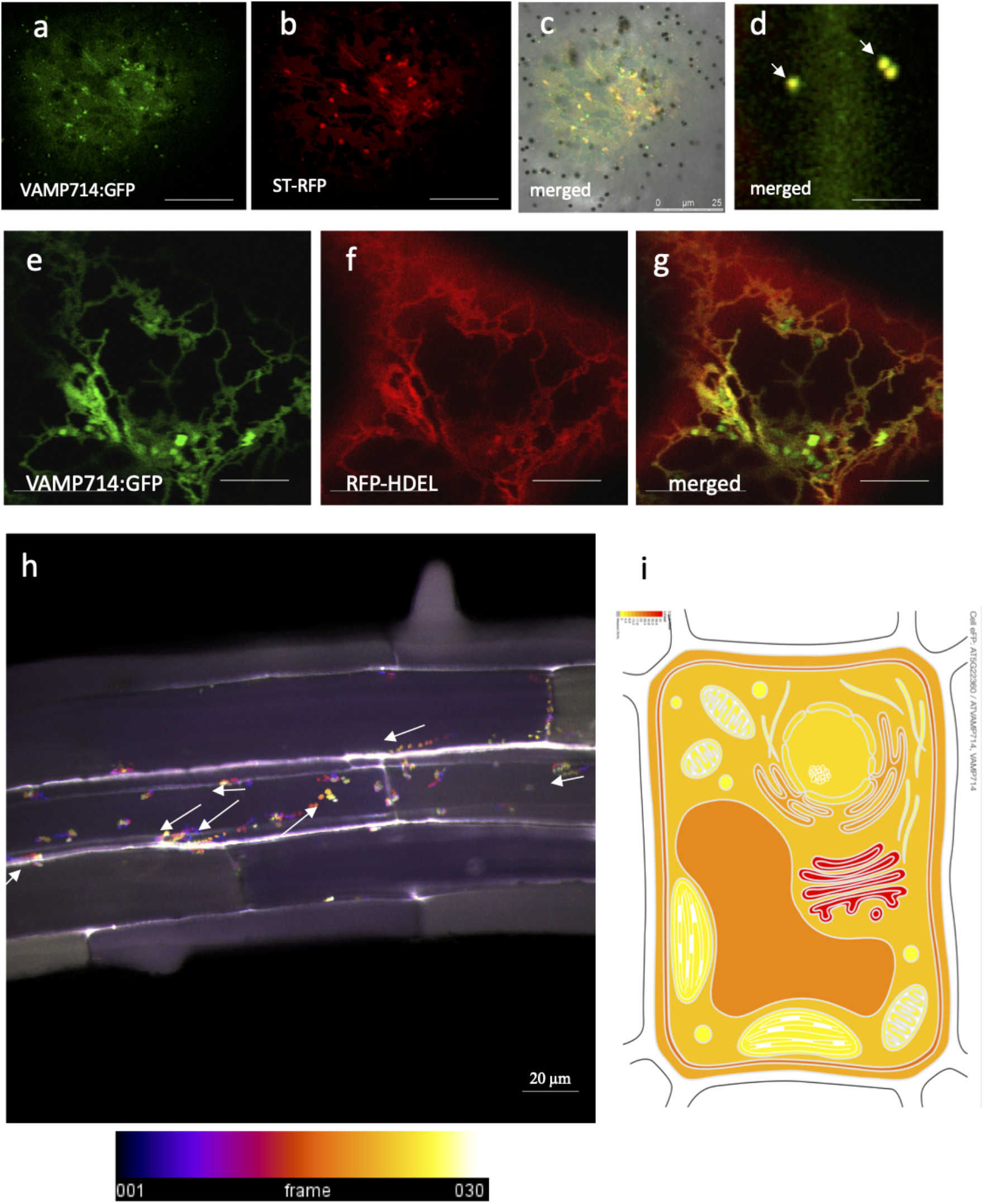
VAMP714 fusion proteins localize to vesicles. (a-d) Transient expression and localization of VAMP714:GFP (a) and the Golgi membrane marker ST–RFP (b), showing co-localization in merged images (c,d; arrowheads indicate individual vesicles showing co-localization). Bars = 25 μm (a-c), 10 μm (d). (e-g) Transient expression and localization of VAMP714:GFP (e) and the ER membrane marker RFP-DEL (f), showing some co-localization in merged images (g). Bars = 15 μm (h) Still image captured from a video (see Supplementary Video 1 for video sequence) with temporal-color code tracking of VAMP714:mCherry-positive vesicle movement. White arrows indicate the direction of vesicle transport to the plasmamembrane. Bar = 20 μm. (i) Heat map of predicted VAMP714 subcellular location, using online tool at http://bar.utoronto.ca/eplant/, showing highest levels (red) at the Golgi, ER and plasmamembrane.

The spatial expression pattern of the *VAMP714* gene was examined in seedlings expressing a promoter reporter fusion (*proVAMP714::GUS*) using histochemical localization of GUS activity (nine independent transgenic lines showed similar patterns of GUS activity; representative images are shown in Fig. S2a-e). GUS activity was detected in roots, and most strongly in vascular tissues of primary and lateral roots, though also at lower levels in the root cortex and in the QC; and at relatively low levels in cotyledon veins, but not in leaf. This expression pattern is consistent with data from the analysis of the transcriptomes of individual root cell types in day 6 seedlings (Birnbaum *et al*., 2003; Nawy *et al*., 2003; and visualized at www.bar.utoronto.ca; Fig. S2f).

Since primary and lateral root growth, correct columella patterning, and shoot branching control are dependent on correct auxin transport and/or signal transduction, and *VAMP714* is expressed in roots and vascular tissues that have relatively high auxin responses (Perret *et al*., 2009; Sabatini *et al*., 1999), these observations led us to hypothesize a role for *VAMP714* in auxin signalling.

To investigate auxin responses in mutant and overexpressing plants, we measured the transcription of the auxin-regulated genes *IAA1* and *IAA2* (Hagen & Guilfoyle, 2002) at 7 dpg by qRT-PCR. The transcript levels of both genes were found to be reduced compared to wildtype in *vamp714* insertional mutants, dominant negative mutants and also in *VAMP714* overexpressors (Fig. 5a). Histochemical analysis of the auxin reporter genes *IAA2::GUS* (Swarup *et al*., 2001) and *DR5::GFP* (Sabatini *et al*., 1999) revealed altered expression patterns in the *vamp714* mutants and overexpressor (Fig. 5b,c). Compared to wildtype, *IAA2::GUS* staining is distally shifted to the disorganized columella of both *vamp714* mutant and overexpressing seedlings (Fig. 5b); while *DR5::GFP*, which is mainly detected in the quiescent centre and columella in the wildtype, exhibits a broadly similar spatial pattern in the roots of the mutants and overexpressers to wildtype but reveals the disorganized cellular patterning of the mutants and overexpressers (Fig. 5c). These data are indicative of incorrect auxin distribution or auxin content in the root tip and demonstrate that wildtype *VAMP714* expression is required for correct auxin distribution and responses. Gravitropism is an auxin-mediated response and linked to correct function of starch-containing columella cells (Wolverton *et al*., 2011), and in gravitropism assays, only 10% of *vamp714* roots showed a true gravitropic response, compared to 85% of wildtype roots at 24 h (n = 20; Fig. 5d,e). This further supports a role for VAMP714 in auxin-mediated processes.

**Figure 5.**
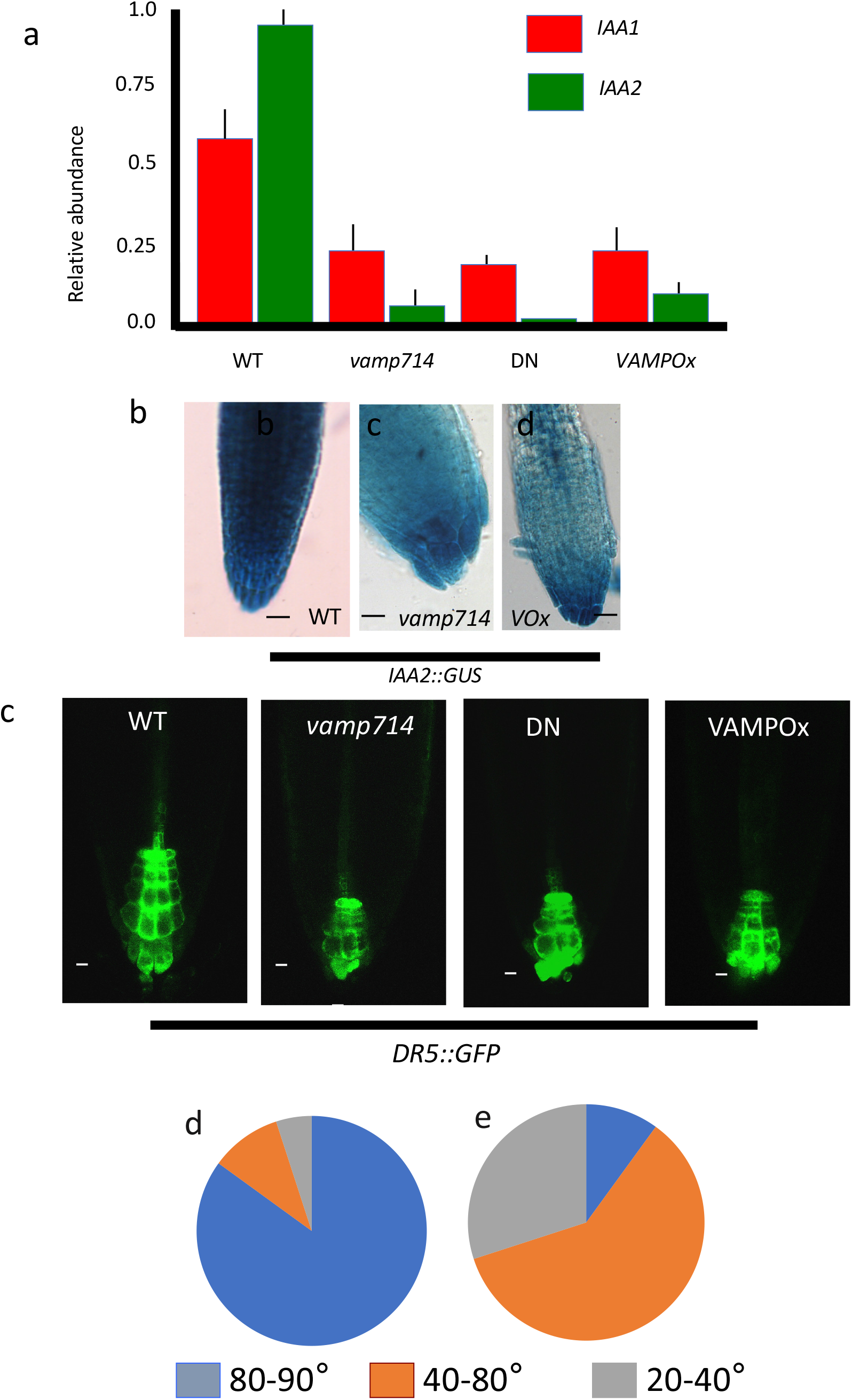
VAMP714 is required for correct auxin-mediated gene expression and root gravitropism. (a) qRT-PCR analysis of mRNA abundance for the auxin-inducible *IAA1* and *IAA2* genes in wildtype (WT), *vamp714* mutant, VAMP714 dominant negative mutant (DN) and transgenic overexpressing (VAMP Ox) seedlings at 7 dpg, compared to *ACTIN2* expression. (b-d) *proIAA2::GUS* reporter activity in wildtype (WT, b), *vamp714* mutant (c) and transgenic overexpressing (VOX, d) roots at 7 dpg. Bars = 10 μm. (e, f) *proDR5::GFP* expression in wildtype (WT), *vamp714* mutant, transgenic dominant negative VAMP714 mutant (DN) and transgenic overexpressing (VAMPOx) roots at 7 dpg. Bars = 10 μm. (d,e) Diagrammatic representation of the gravitropic responses of wildtype (d) and *vamp714* mutants (e) at 24 h after shifting the vertical axis by 90°. The pie-charts indicate the proportion of seedlings showing bending responses at between 80° to 90° from horizontal (blue), 40° to 80° from horizontal (orange) and 20° to 40° from horizontal (grey).

Given that VAMP714 is required for correct auxin patterning and responses, we considered that it might itself be activated in response to auxin, since for example auxin promotes *PIN* gene expression and PIN protein localization (Paciorek *et al*., 2005; Heisler *et al*., 2005). To investigate this hypothesis, wildtype seedlings were transferred to medium containing 10 μM IAA and the steady state transcript levels of *VAMP714* were measured after 12, 24 and 36 h. The auxin treatment increased relative transcript abundance for the *VAMP714* gene ca. 3 fold by 24 h after treatment, compared to an *ACTIN2* internal control gene (Fig. 6a).

**Figure 6.**
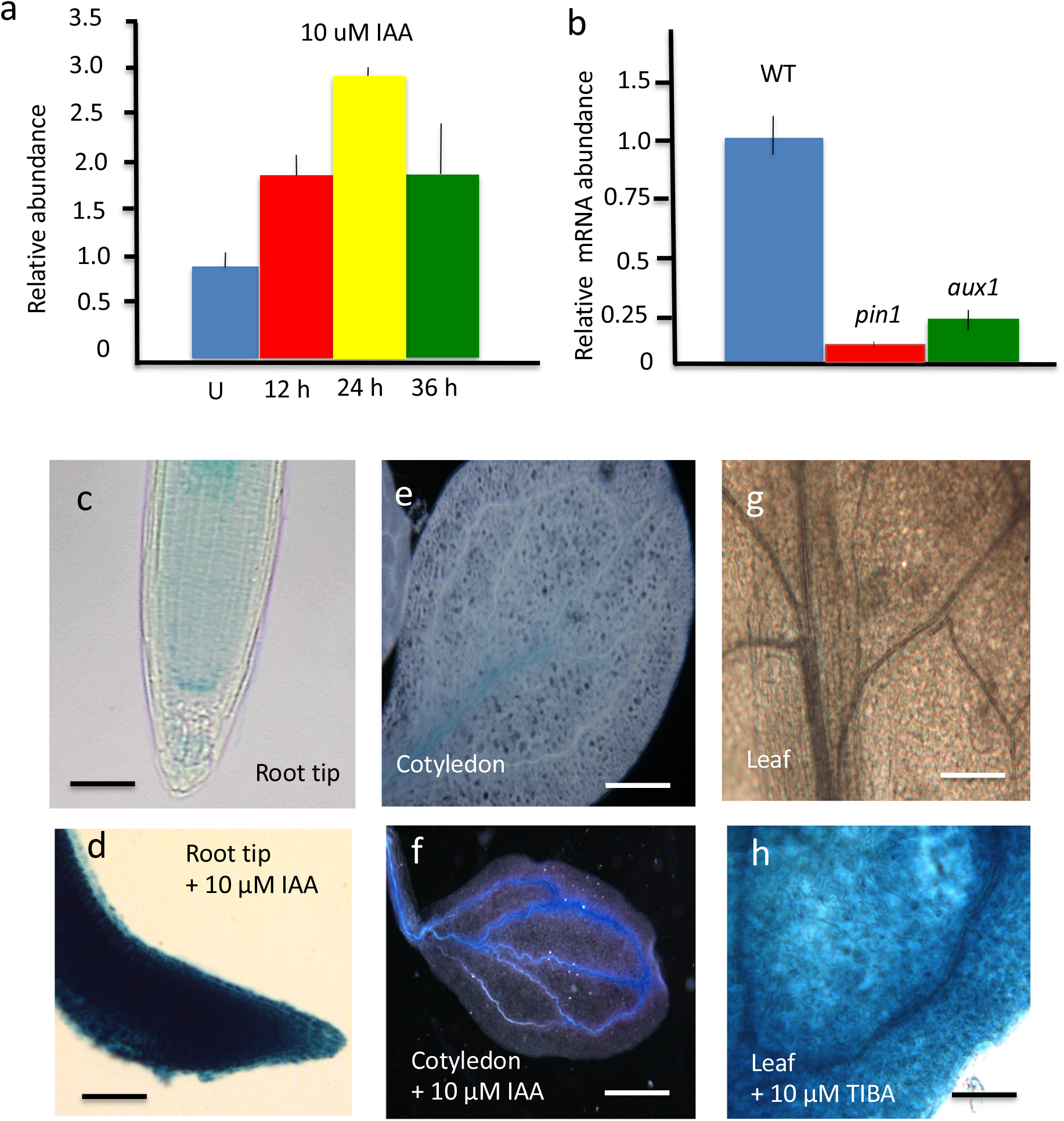
VAMP714 gene is auxin-regulated. (a) qRT-PCR analysis of mRNA abundance of *VAMP714* seedlings either untreated (U) or treated with 100 μM IAA for 0, 12, 24 and 36 h. Expression levels are relative to *ACTIN2* expression. Data represent means of 3 biological replicates ± SD. (b) qRT-PCR analysis of mRNA abundance of *VAMP714* in wildtype, *pin1* and *aux1* mutant seedlings at 7 dpg. Expression levels are relative to *ACTIN2* expression. Data represent means of 3 biological replicates ± SD. (c,d) Primary root tip of 7 dpg seedling either untreated (c) or treated with 10 μM IAA for 5 days (c), bars = 25 μm. (e,f) Cotyledon of seedlings either untreated (e) or treated with 10 μM IAA for 5 days (f), bars = 25 μm (e), 40 μm (f). (g,h) Leaf of seedling either untreated (g) or treated with 10 μM TIBA for 5 days (h), bars = 25 μm (g), 30 μm (h)

To study the dependence of *VAMP714* expression on correct auxin transport and signalling *in planta*, we compared *VAMP714* expression in *pin1* and *aux1* mutants with wildtype. The *pin1* and *aux1* mutants exhibit reduced polar auxin transport (Okada *et al*., 1991; Bennett *et al*., 1996). In both mutants, the level of *VAMP714* mRNA was significantly reduced compared to wildtype (Fig. 6b).

While exogenous cytokinin and ACC had no detectable effect on *proVAMP714::GUS* expression (data not shown), exogenous auxin (10 μM IAA) induced strong GUS activity in root tips (Fig. 6c,d), and in cotyledon vascular tissues (Fig. 6e,f). 10 μM TIBA treatment, which induces the accumulation of auxin in aerial tissues by inhibition of polar auxin transport, led to an activation of GUS activity in the young leaf (Fig. 6g,h). Consistent with its auxin inducibility, sequence analysis of a 2 kb promoter region upstream of the *VAMP714* gene revealed the presence of an auxin-response element (AuxRE) motif (TGTCTC) (Sabatini *et al*., 1996) at position −1346, though the functionality of this element was not tested experimentally. The observed auxin-inducibility of expression is consistent with *VAMP714* transcription in vascular and QC cells, which contain relatively high concentrations of auxin (Sabatini *et al*., 1996; Dengler, 2001).

In view of the diverse auxin signalling-related defects in *vamp714* mutants and overexpressers, and the prospective role for VAMP714 in membrane vesicles, we investigated a possible role for VAMP714 in PIN localization and polar auxin transport. In wildtype cells, PIN1:GFP was localized as expected to the basal membrane of the cells in the central cylinder (Fig. 7a), and PIN2:GFP was localized to the apical membrane of the cells in the root cortex and epidermis (Fig. 7b), as expected. Both PIN1 and PIN2 were less concentrated at the plasmamembrane in mutant, dominant negative and overexpressing plants (Fig. 7c). The reduction of PIN localisation at the plasmamembrane was accompanied by higher protein levels in the cytoplasm, resulting in lower values of membrane:cytoplasm ratios of PIN1 and PIN2 in the null mutant (Fig. 7d). In transgenic plants expressing *proVAMP714::VAMP714:CFP*, both PIN1:GFP and VAMP714:CFP, and PIN2:GFP and VAMP714:CFP, co-localize at the plasmamembrane, though less clearly for PIN2 than for PIN1 (Fig. 7a,b). This may be linked to the stronger expression of the *VAMP714* gene in vascular tissues, where PIN1 is strongly expressed, while PIN2 is localized to epidermal and cortical cell layers.

**Figure 7.**
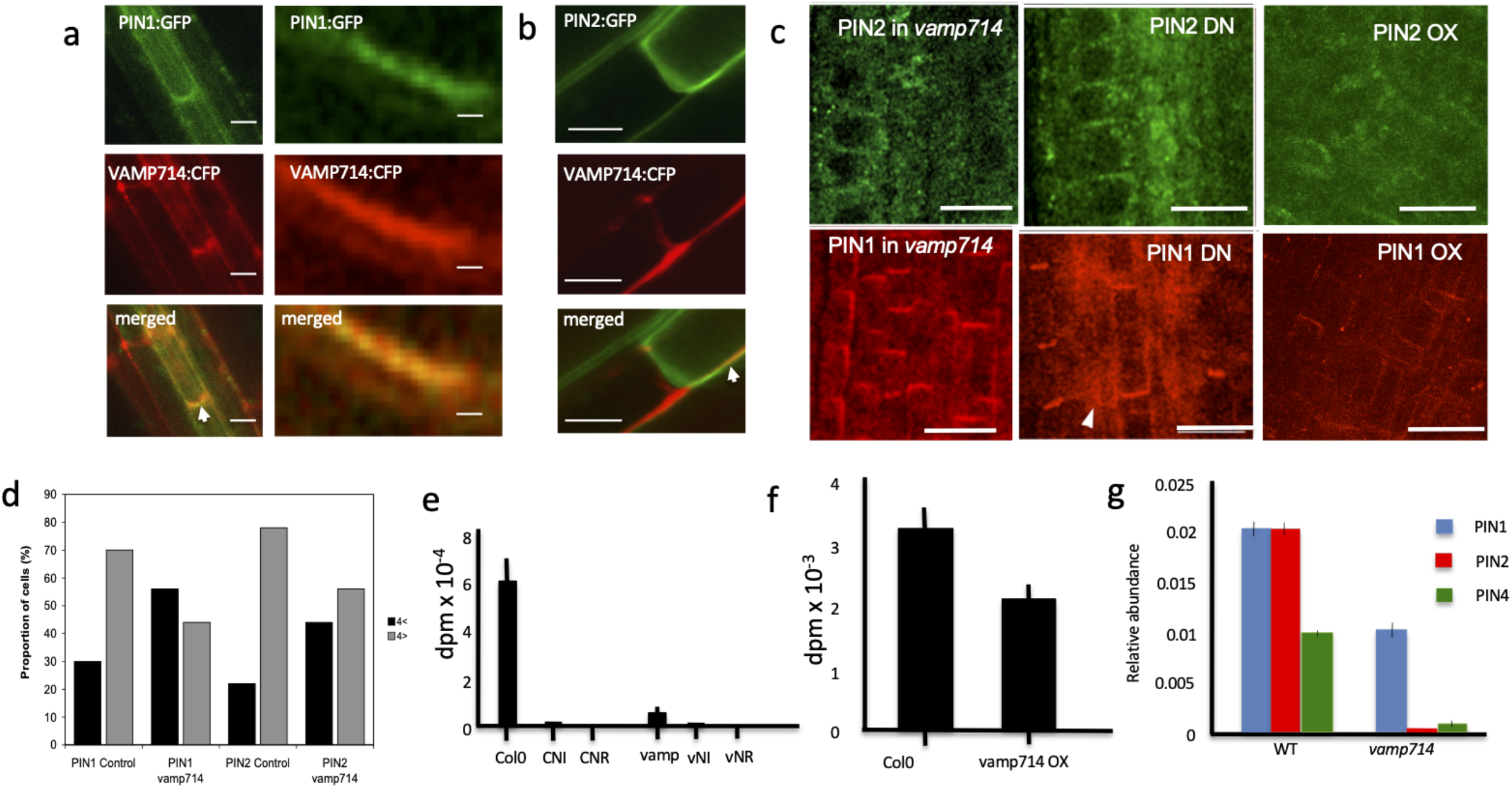
VAMP714 is required for correct PIN protein localization and polar auxin transport. (a) proPIN1::PIN1:GFP and (b) proPIN2::PIN2:GFP localization (upper panels) and co-localization with proVAMP714::VAMP714:CFP (lower two panels) at the basal plasmamembrane of root vascular cells in transgenic plants. Arrowheads highlight co-localization in merged images. (a) left panels: bars = 10 μm; (a) right panels: bars = 1 μm; (b) bars = 10 μm. (c) PIN1 and PIN2 localization in seedling roots of *vamp714* mutants (left two panels), dominant negative (DN, centre two panels) and *pro35S::VAMP714* overexpressers (OX, right two panels) at 7 dpg. Arrowhead exemplifies disrupted PIN localization. Bars = 20 μm. (d) Quantification of PIN1 and PIN2 distribution in wildtype and *vamp714* mutant cells, showing the proportion of cells with relatively strong fluorescence signal at the plasma membrane (four-fold above the cytoplasmic signal and above; grey bars) versus relatively weak signal at the plasmamembrane (less then four-fold above the cytoplasm level; black bars) for wildtype (control) and mutant (*vamp714*). The mutant exhibits a lower percentage of cells showing the fluoresence signal for both PIN1 and PIN2 at the plasmamembrane. (e, f) Polar auxin transport measurements in (e) inflorescence stems of *vamp414* mutants and (f) roots of *pro35S::VAMP714* misexpressors. (e) Col-0 indicates auxin transport in the wildtype control; CNI is the non-inverted wildtype control (the stem was not inverted, so that the basal region was exposed to the ^3^H-IAA); CNR is the wildtype control in non-radiactive medium; vamp indicates auxin transport in the *vamp714* mutant; vNI is the non-inverted *vamp714* mutant; vNR is *vamp714* mutant incubated in non-radioactive medium. Data represent the means of 5 independent assays ± SD. (f) Auxin transport assays in wildtype (wt) and transgenic *pro35S::VAMP714-overexpressing* (VAMP Ox) roots. Data represent the means of 5 independent assays ± SD. (g) qRT-PCR analysis of mRNA abundance of *PIN1, PIN2* and *PIN4* genes in wildtype (WT) and *vamp714* mutant seedlings, relative to *ACTIN2* expression. Error bars represent means ± SD of 3 biological replicates.

To examine the effect of aberrant PIN localisation on auxin transport, we used a 3 [^3^H]-IAA transport assay. The rate of auxin transport was significantly reduced in both hypocotyl (Fig. 7e) and root (Fig. 7f) of the PIN localization-defective *VAMP714* misexpressors, compared with wildtype controls. Given the proposed regulatory loop in which auxin promotes *PIN* gene expression which then regulates directional auxin transport (Grieneisen *et al*., 2007), we hypothesized that the levels of *PIN* gene expression in the loss-of-function *vamp714* might also be reduced. Consistent with this hypothesis, the transcription of *PIN1*, *PIN2* and *PIN4* genes was reduced in the *vamp714* loss-of-function mutant (Fig. 7g).

The PIN proteins are dynamically regulated in their subcellular localization via the endsome recycling pathway (Geldner *et al*., 2001), and we hypothesized that VAMP714-associated vesicles may also be subject to endosome recycling. This recycling is inhibited by both the vesicle-trafficking inhibitor brefeldin A (BFA), leading to the intracellular accumulation of BFA bodies, and by the actin depolymerizing agent latrunculin B (LatB) (Geldner *et al*., 2001). To determine whether VAMP714 is also subject to actin-dependent endosome recycling, we treated *proVAMP714::VAMP714:mCherry* seedlings with 50 μM BFA or 20 μM LatB, and monitored the formation of VAMP-positive BFA bodies in root cells. We also treated *proPIN1::PIN1:GFP* and *proPIN2::PIN2:GFP* seedlings with 50 μM BFA as positive controls. The VAMP714:mCherry fusion protein was demonstrated to be biologically functional, as shown by transgenic complementation of the *vamp714* loss-of-function mutant (Fig. 1c). We found that VAMP714, PIN1 and PIN2 exhibit the same BFA body formation, which can be washed out (Fig. 8a), indicative of endosome recycling between BFA compartments and the plasma membrane. We also found that LatB caused intracellular accumulation of VAMP714 vesicles (Fig. 8b). This suggests that VAMP714 forms part of both the exocytic vesicle trafficking pathway from the ER/Golgi and the actin-dependent endocytic recycling pathway, which together regulate PIN protein concentrations at the plasma membrane. Relatively high intracellular levels of both PIN1:GFP and PIN2:GFP, and some intracellular PIN1:GFP-positive vesicle-like structures are seen in the *vamp714* mutant, DN and overexpressers compared with wildtype (Fig. 8c), broadly consistent with the observations for PIN immunolocalization (Fig. 7c) and indicative of a requirement of VAMP714 for polar PIN localization.

**Figure 8.**
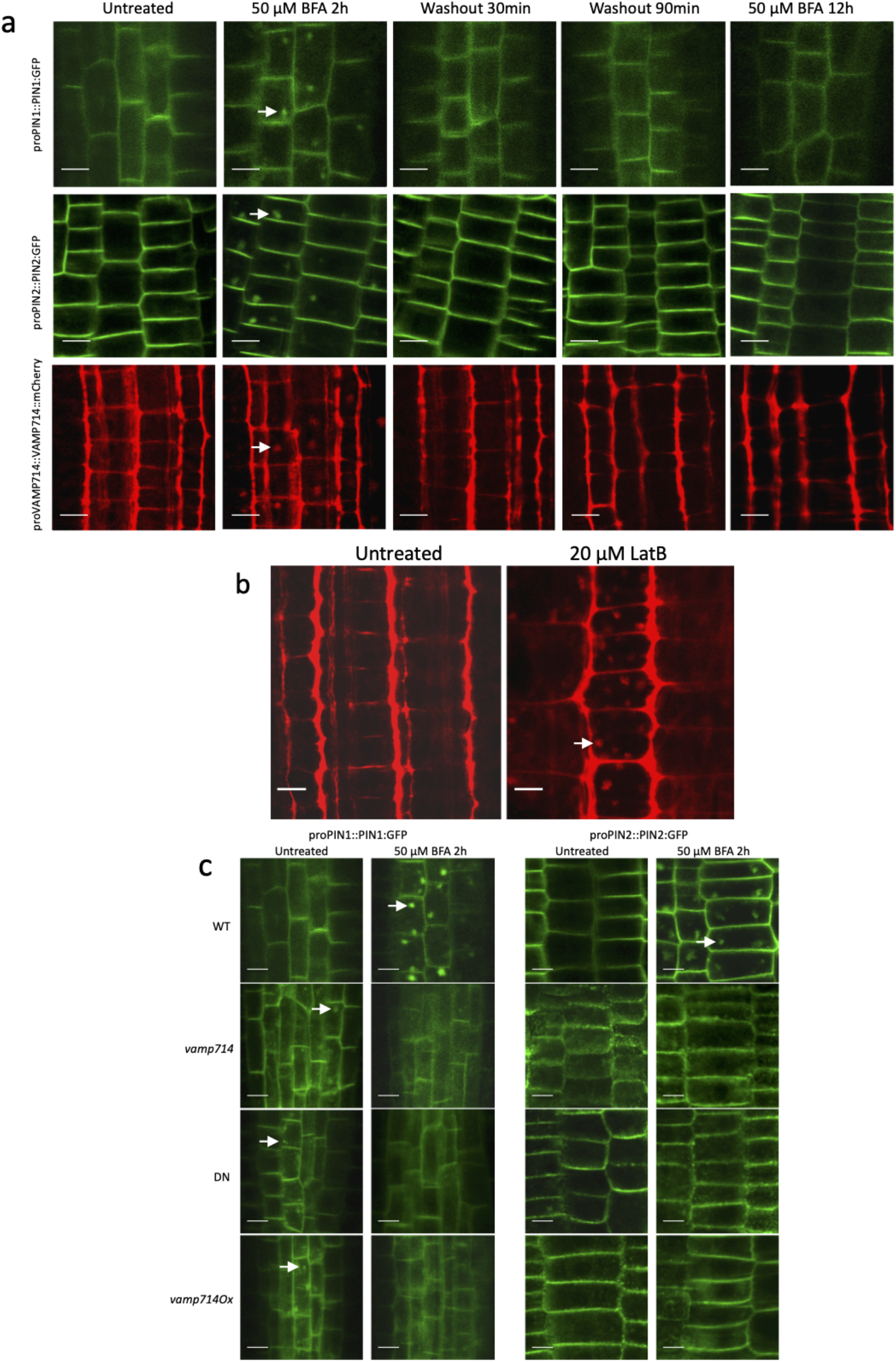
VAMP714, PIN1 and PIN2 exhibit same endosome recycling from BFA compartments to plasma membrane and actin requirements for polar VAMP714 targeting. (a) Seedling roots expressing proPIN1::PIN1:GFP, proPIN2::PIN2:GFP and proVAMP714::VAMP714:mCherry were imaged before and after 2 h of treatment with 50 μM brefeldin A (BFA), 30 and 90 min after BFA washout, and after a prolonged 12 h treatment with BFA. The localization of PIN:GFP proteins and VAMP714:mCherry proteins in the plasma membrane was re-established by 30 min after washout in the wild type. Arrows indicate BFA bodies. Bars = 10 μm. (b) Seedlings expressing proVAMP714::VAMP714:mCherry were imaged before and after 3 h of treatment with 20 μM latrunculin B, revealing sensitivity to actin depolymerization. Arrow indicates intracellular vesicle accumulation. Bars = 10 μm. (c) wildtype (WT), *vamp714* mutant, and VAMP714 dominant negative mutant (DN) seedlings expressing either proPIN1::PIN1:GFP or proPIN2::PIN2:GFP imaged before treatment with 50 μM brefeldin A (BFA; untreated, left panels) and after 2 h of BFA treatment (right panels). WT seedlings exhibited PIN:GFP internalization in BFA compartments, whereas *vamp714* mutant and dominant negative mutant seedlings showed no PIN accumulation in BFA bodies. Arrows indicate intracellular accumulation of PIN. Bars = 10 μm.

We therefore also investigated the role of VAMP714 in endocytic recycling. We monitored PIN1 and PIN2 recycling in the *vamp714* loss-of-function and dominant negative mutants in the presence of 50 μM BFA. We found that PIN accumulation in BFA bodies does not occur in either mutant background (Fig. 8c). This suggests that VAMP714 is required for PIN endosome recycling.

## Discussion

Auxin homeostasis, transport and signalling each play major roles in multiple developmental pathways in plants, and directional transport is key to establishing functional concentration gradients of auxin that mediate the control of cell identity, tropic growth and the nature of interactions with other hormones to elicit specific responses (Benjamins & Scheres, 2008). Directional transport of auxin is principally mediated by PIN protein family members, some of which become localized to specific faces of the cell plasma membrane; and expression of *PIN* genes appears to reflect local auxin concentrations, reflective of a feedback regulatory system (Omelyanchuk *et al*., 2016). PIN localization involves an actin-mediated recyling between the plasma membrane and endosomes, providing a mechanism for rapid changes in the placement of these transporters. It is now well established that ARF GEF- and Rab5 GTPase-dependent recycling is critical for PIN localization, and this process is itself inhibited by BFA (an ARF GEF inhibitor) and modulated by auxin (Steinmann *et al*., 1999; Geldner *et al*., 2001; Kleine-Vehn *et al*., 2008; Kitakura *et al*., 2011). Less clear have been the mechanisms regulating the exocytic delivery of PIN proteins from the ER/Golgi to the plasma membrane. We show in this paper that the Arabidopsis R-SNARE VAMP714 is required for correct PIN localization, likely via both the exocytic and endosome recycling pathways.

SNAREs have been classified as vesicle-associated (v-SNAREs) and target membrane-associated SNAREs (t-SNAREs) (Sollner *et al*., 1993), though under a structural classification they can be grouped as Q- and R-SNAREs, owing to the occurrence of either a conserved glutamine or arginine residue in the centre of the SNARE domain (Fasshauer *et al*., 1998). Generally, t-SNAREs correspond to Q-SNAREs, and v-SNAREs correspond to R-SNAREs. There are more than 60 SNARE protein-encoding genes represented in the Arabidopsis genome (Uemura *et al*., 2004; Lipka *et al*., 2007; Sanderfoot, 2007), but there is limited information available on the roles of SNARE proteins from genetic studies in plants, most likely because of a lack of loss-of-function phenotypes due to functional redundancy between related family members. For example, redundancy has been demonstrated between VTI11 and VTI12 (Kato *et al*., 2002; Surpin *et al*., 2003), SYP121 and SYP122 (Assaad *et al*., 2004; Zhang *et al*., 2008), and VAMP721 and VAMP722 (Kwon *et al*., 2008).

The animal VAMP synaptobrevin has been implicated in linking synaptic vesicles to the plasma membrane (Walch-Solimena *et al*., 1993; Bonifacino & Glick, 2004). It is proposed that R-SNAREs may play a key role in determining specificity in vesicle budding, and an important mechanism for SNARE localization is interaction with vesicle coats. For example, it has been shown that R-SNAREs may be components of the COPII vesicles that are involved in ER-Golgi transport (Springer & Schekman, 1998), and that R-SNAREs must be packaged into COPI vesicles during retrieval from the Golgi (Rein *et al*., 2002).

The data presented in this paper provides new information on both the role of the R-SNARE VAMP714 and the molecular components required for the control of auxin transport and auxin-mediated responses via PIN protein expression, recycling and localization. We propose a model in which the correct delivery of PIN proteins from the ER/Golgi to the plasma membrane is via a VAMP714-associated compartment, that is a necessary precursor to the endocytic recycling that provides dynamical control over the level and site of PIN protein localization (Fig. S3). This in turn regulates the direction and rate of auxin efflux. We show that VAMP714 is required for a range of correct auxin responses, including auxin-mediated gene expression, root gravitropism, root cell patterning and shoot branching.

This model is supported by the co-localization of VAMP714 and PIN proteins at the plasma membrane; the accumulation of PIN proteins in the cytoplasm in *vamp714* loss-of-function and dominant negative mutants and VAMP714 misexpressers; and the requirement for wildtype levels of expression of *VAMP714* for BFA body formation (i.e. endosome recycling). Significantly, SNARES and Rab GTPases have been demonstrated to interact functionally to promote vesicle fusion at the endosome, and act coordinately to increase the specificity and efficiency of membrane fusion (Ohya *et al*., 2009; Ebine *et al*., 2011). Mechanistically VAMP714 may interact with the RAB5 GTPase complex known to participate in PIN recycling at the endosome (Dhonukshe *et al*., 2008), following its exocytic transport of PINs, and this possibility is the subject of further studies. It is also currently unclear whether VAMP714 is involved in transcytosis to modulate PIN localization; and whether it is required for PIN-specific or more general transport of plasma membrane proteins.

In the classical canalization hypothesis pioneered by Sachs (Sachs, 1981), auxin itself promotes its own transport system, leading to directional flow through tissues and subsequent establishment of cell polarity and differentiation. Consistent with this hypothesis, the auxin-mediated transcriptional activation of the *VAMP714* gene would allow the activation of a pathway essential for polar auxin transport by promoting correct PIN protein localization at the plasma membrane. Integrated in this mechanism would be auxin-mediated transcriptional effects on *PIN* genes (Heisler *et al*., 2005) and the effect of auxin on PIN endocytosis (Paciorek *et al*., 2005). A role for VAMP714 in the (probably indirect) maintenance of *PIN* gene expression is also indicated. Our studies demonstrate that R-SNARE-dependent exocytosis is essential for the auxin transport and downstream signalling pathways that are required for the control of cell polarity, tropic growth and morphogenesis in plants.

## Acknowledgements

KL acknowledges The Biotechnology and Biological Sciences Research Council (BBS/B/0773X) and Durham University for funding. XG thanks the China Scholarship Council for funding to support her studies at Durham University.

## Author contributions

KL, SAC and JFT devised the project; XG, KF and SAC carried out the experimental work; KL, JFT, PJH and GG supervised the work; KL drafted the manuscript; all authors edited the manuscript.

## Data availability statement

All materials and data described in this papaer are available to readers from the corresponding author, upon reasonable request.

## Supporting Information

**Figure S1. Construction and analysis of dominant negative VAMP714 plants.**

(a) Domain structure of AtVAMP714, showing sites of primers used to contruct a negative dominant protein gene.

(b) AtVAMP714 Domain DNA sequence and primers used.

(c) Domain amino acid sequences.

(d) *AtVAMP714* gene expression in dominant negative transgenics.

(e) Phenotypes of dominant negative and wildtype plants.

**Figure S2. VAMP714 expression in Arabidopsis.**

(a-e) *proVAMP714::GUS* is expressed in vascular tissues.

(a) Whole seedling, 4 dpg, bar = 1 mm.

(b) Seedling root, 4 dpg with transverse section in mature region of root, showing GUS activity in the stele (b’), bar = 1 mm.

(c) Cotyledon at 4 dpg, bar = 1 mm.

(d) root-hypocotyl junction at 4 dpg, bar = 1 mm.

(e) primary root tip, 4 dpg, bar = 100 μm.

(f) Expression heat map of VAMP714 gene in primary root of Arabidopsis.

Visualized using online tool at http://bar.utoronto.ca/eplant/. Red denotes high expression, yellow denotes low expression.

**Figure S3. Proposed model for the role for VAMP714 in exocytosis and endosomal cycling of PIN proteins.**

Our data show that VAMP714:mCherry-positive vesicles (purple) move towards the plasma membrane, and co-localize at the plasma membrane (PM) with PIN proteins. VAMP714 also accumulates in BFA bodies and in aggregates following latrunculin B treatment, in the same manner as PIN proteins, both processes being part of endosome recycling (green). VAMP714 is also required for PIN1-positive BFA body formation. It is therefore proposed that VAMP714 is required for both exocytosis of PIN vesicles to the plasma membrane and for PIN cycling between the plasma membrane and endosomes, a process sensitive to BFA and latrunculin B in Arabidopsis.

**Video S1. VAMP714 localizes to the plasmamembrane via vesicle trafficking.**

Video showing time series of VAMP714:mCherry expression (30 images were captured over 10 minutes).

