## Supplementary material for "The Arabidopsis R-SNARE VAMP714 is essential for polarization of PIN proteins in the establishment and maintenance of auxin gradients": Suppl. Figs 1-3

**Supplementary Figure 1.**

**a. Domain structure of AtVAMP714.**

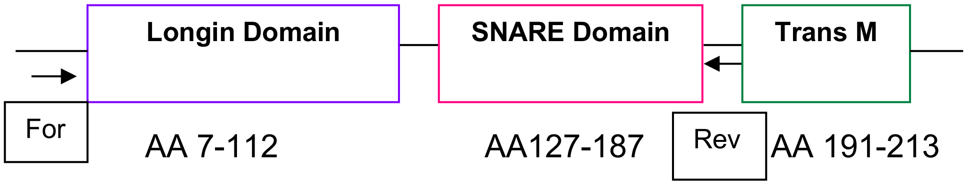


The three main domains of AtVAMP714 (Longin domain, SNARE domain and Transmembrane domain (Trans M) with the length of the amino acids. For and Rev indicates the position of forward and reverse primers designed for amplifying the Longin and SNARE domains, which were cloned into the Gateway vector pMDC43, under the transcriptional control of the CaMV35S gene promoter.

**b. AtVAMP714 Domain DNA sequence and primers used.**

*For* primer (Pink) and *Reverse* primer (Blue) were used to amplify the Longin and SNARE domains to create VAMP714 dominant negative transgenic plants.

```
At5g22360/ VAMP714
Longin Domain
SNARE Domain
Trans-membrane Domain
For primer
Reverse primer

   1  CATTTTTATA CTCTGTTCTG ATCGCAGCAA AGCCGACGTT GAACTTTCTC
  51  GCCGCCGGAG CGCGTGATCT CCACTCTCTG TCATCGAATC ACTCTAATTG
 101  AAGATTCTCC GATGGCGATT GTCTATGCTG TTGTAGCGAG AGGTACCGTG
 151  GTATTAGCTG AATTCAGCGC CGTTACGGGA AACACAGGCG CCGTGGTGCG
 201  ACGGATCCTC GAGAAGCTTT CACCGGAAAT CTCCGATGAA AGACTTTGTT
 251  TCTCTCAAGA TCGTTATATC TTCCATATTC TTAGATCTGA TGGTCTTACC
 301  TTTCTCTGTA TGGCCAATGA TACCTTTGGA AGGAGGGTTC CATTTTCGTA
 351  TTTGGAAGAG ATTCATATGA GATTCATGAA AAACTATGGC AAAGTGGCTC
 401  ATAATGCTCC AGCTTATGCA ATGAATGATG AATTCTCAAG GGTTTTGCAT
 451  CAGCAGATGG AGTTCTTCTC TAGTAATCCT AGTGTTGATA CTCTCAATCG
 501  TGTTAGAGGA GAAGTCAGTG AGATTCGATC GGTCATGGTA GAGAACATTG
 551  AGAAGATAAT GGAAAGAGGT GATAGGATTG AGCTTCTTGT TGATAAAACA
 601  GCAACAATGC AAGATAGCTC GTTTCACTTC AGGAAGCAAT CTAAGCGCCT
 651  TCGCCGAGCT CTTTGGATGA AAAATGCTAA GCTCCTGGTC TTGTTGACAT
 701  GCTTGATAGT TTTCTTGCTG TACATAATAA TCGCATCTTT CTGCGGAGGA
 751  ATCACTTTAC CATCATGCAG ATCTTAAAAT CTGGCGGCCT TATCTAAGGT
 801  ATACTGAAAC GGACCACTGT TTTTGTAAAT CAACTCAGTC GCATCATTTC
 851  GATTTGAAGC CTTGGTTTTG TCATGAAAAT GACTGTGAGT TTGAAGTTAC
 901  ATGTCATGGT CCTCCTTGCT TATGTAACTC TTGTAAATGT CAAAATCAAA
 951  ATGATACAGA GGTTCATTGA ACTCTTGCCT TGTCTTTTAG ATTTAGTCAC
1001  GTGAGTGAGT TTGTTCTCTG TATTTCCAAA ACTTTATCCG CTGTGTCTTT
1051  ATAATTTTTG CATTTGCAAT GTACATCACA TATTCGTTAT TTTTGTTGTT
1101  AAAATTACTT CAGTTTTCAT CTTTGTTTAA TAACATTTCT GATTCACAAA
1151  TA
```

**c. Domain amino acid sequences, highlighted in different colours (Longin, SNARE and Transmembrane).**

| | | | | | | | | | | | | | | | | | | |
|---|---|---|---|---|---|---|---|---|---|---|---|---|---|---|---|---|---|---|
| | Met | Ala | Ile | Val | Tyr | Ala | Val | Val | Ala | Arg | Gly | Thr | Val | Val | Leu | Ala | Glu | 17 |
| 112 | ATG | GCG | ATT | GTC | TAT | GCT | GTT | GTA | GCG | AGA | GGT | ACC | GTG | GTA | TTA | GCT | GAA | |
| | Phe | Ser | Ala | Val | Thr | Gly | Asn | Thr | Gly | Ala | Val | Val | Arg | Arg | Ile | Leu | Glu | 34 |
| 163 | TTC | AGC | GCC | GTT | ACG | GGA | AAC | ACA | GGC | GCC | GTG | GTG | CGA | CGG | ATC | CTC | GAG | |
| | Lys | Leu | Ser | Pro | Glu | Ile | Ser | Asp | Glu | Arg | Leu | Cys | Phe | Ser | Gln | Asp | Arg | 51 |
| 214 | AAG | CTT | TCA | CCG | GAA | ATC | TCC | GAT | GAA | AGA | CTT | TGT | TTC | TCT | CAA | GAT | CGT | |
| | Tyr | Ile | Phe | His | Ile | Leu | Arg | Ser | Asp | Gly | Leu | Thr | Phe | Leu | Cys | Met | Ala | 68 |
| 265 | TAT | ATC | TTC | CAT | ATT | CTT | AGA | TCT | GAT | GGT | CTT | ACC | TTT | CTC | TGT | ATG | GCC | |
| | Asn | Asp | Thr | Phe | Gly | Arg | Arg | Val | Pro | Phe | Ser | Tyr | Leu | Glu | Glu | Ile | His | 85 |
| 316 | AAT | GAT | ACC | TTT | GGA | AGG | AGG | GTT | CCA | TTT | TCG | TAT | TTG | GAA | GAG | ATT | CAT | |
| | Met | Arg | Phe | Met | Lys | Asn | Tyr | Gly | Lys | Val | Ala | His | Asn | Ala | Pro | Ala | Tyr | 102 |
| 367 | ATG | AGA | TTC | ATG | AAA | AAC | TAT | GGC | AAA | GTG | GCT | CAT | AAT | GCT | CCA | GCT | TAT | |
| | Ala | Met | Asn | Asp | Glu | Phe | Ser | Arg | Val | Leu | His | Gln | Gln | Met | Glu | Phe | Phe | 119 |
| 418 | GCA | ATG | AAT | GAT | GAA | TTC | TCA | AGG | GTT | TTG | CAT | CAG | CAG | ATG | GAG | TTC | TTC | |
| | Ser | Ser | Asn | Pro | Ser | Val | Asp | Thr | Leu | Asn | Arg | Val | Arg | Gly | Glu | Val | Ser | 136 |
| 469 | TCT | AGT | AAT | CCT | AGT | GTT | GAT | ACT | CTC | AAT | CGT | GTT | AGA | GGA | GAA | GTC | AGT | |
| | Glu | Ile | Arg | Ser | Val | Met | Val | Glu | Asn | Ile | Glu | Lys | Ile | Met | Glu | Arg | Gly | 153 |
| 520 | GAG | ATT | CGA | TCG | GTC | ATG | GTA | GAG | AAC | ATT | GAG | AAG | ATA | ATG | GAA | AGA | GGT | |
| | Asp | Arg | Ile | Glu | Leu | Leu | Val | Asp | Lys | Thr | Ala | Thr | Met | Gln | Asp | Ser | Ser | 170 |
| 571 | GAT | AGG | ATT | GAG | CTT | CTT | GTT | GAT | AAA | ACA | GCA | ACA | ATG | CAA | GAT | AGC | TCG | |
| | Phe | His | Phe | Arg | Lys | Gln | Ser | Lys | Arg | Leu | Arg | Arg | Ala | Leu | Trp | Met | Lys | 187 |
| 622 | TTT | CAC | TTC | AGG | AAG | CAA | TCT | AAG | CGC | CTT | CGC | CGA | GCT | CTT | TGG | ATG | AAA | |
| | Asn | Ala | Lys | Leu | Leu | Val | Leu | Leu | Thr | Cys | Leu | Ile | Val | Phe | Leu | Leu | Tyr | 204 |
| 673 | AAT | GCT | AAG | CTC | CTG | GTC | TTG | TTG | ACA | TGC | TTG | ATA | GTT | TTC | TTG | CTG | TAC | |
| | Ile | Ile | Ile | Ala | Ser | Phe | Cys | Gly | Gly | Ile | Thr | Leu | Pro | Ser | Cys | Arg | Ser | 221 |
| 724 | ATA | ATA | ATC | GCA | TCT | TTC | TGC | GGA | GGA | ATC | ACT | TTA | CCA | TCA | TGC | AGA | TCT | |
| | Stop | | | | | | | | | | | | | | | | | 221 |

**d. *AtVAMP714* gene expression in dominant negative transgenics.**

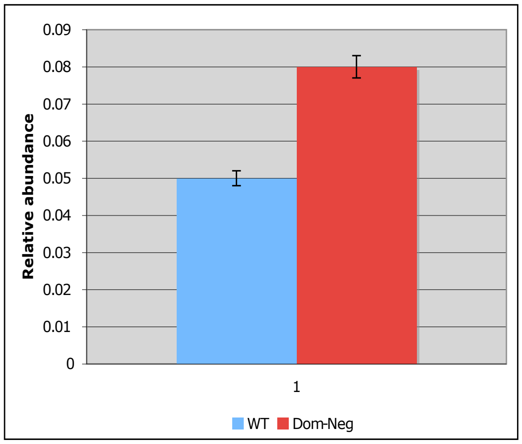


Quantitative RT-PCR analysis of *AtVAMP714* gene expression in pro35S::VAMP714 dominant negative transgenics and Col-0 wild type plants relative to *ACTIN2* at 7 days post germination**.** These data are representative of two independent experiments using biological replicate samples. The error bars represent Standard Errors of the mean of three technical replicates. Y axis represents the relative abundance of the *AtVAMP714* transcript level. PCR primers were designed to detect the truncated form of the dominant negative transcript (Longin and SNARE domains). The relative abundance of the transcript of *AtVAMP714* was higher in dominant negative transgenics than that of Col-0 wild type plants.

**e. Phenotypes of dominant negative and wildtype plants.**

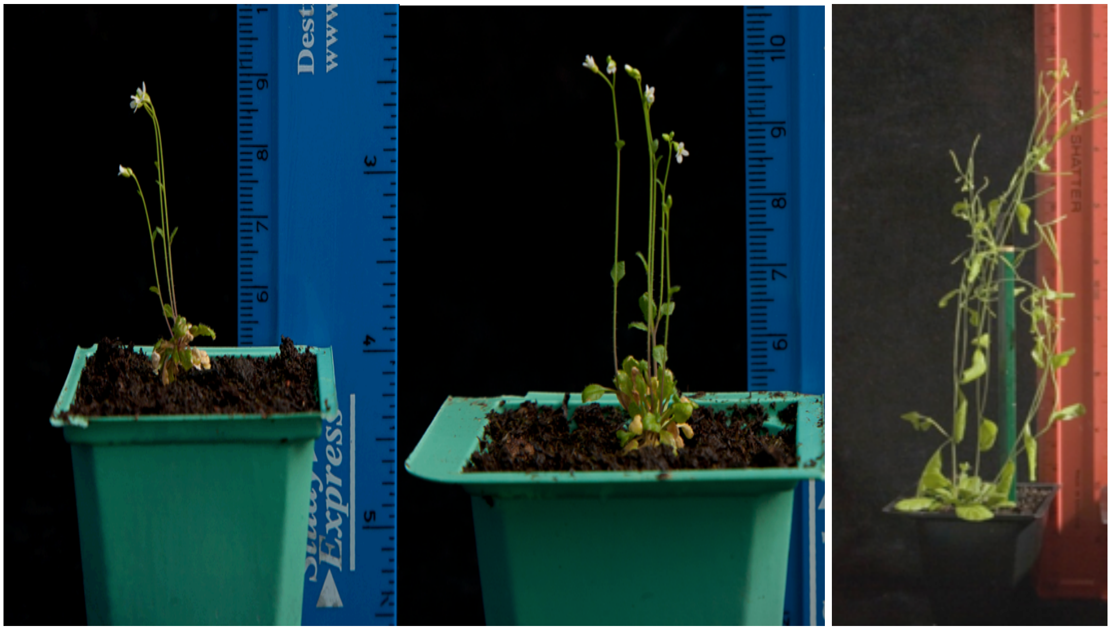


**Left**: Dominant negative transgenic plant at 21 dpg
**Centre**: Dominant negative transgenic plant at 40 dpg
**Right**: Col-0 wildtype plant at 40 dpg

Suppl. Fig 2

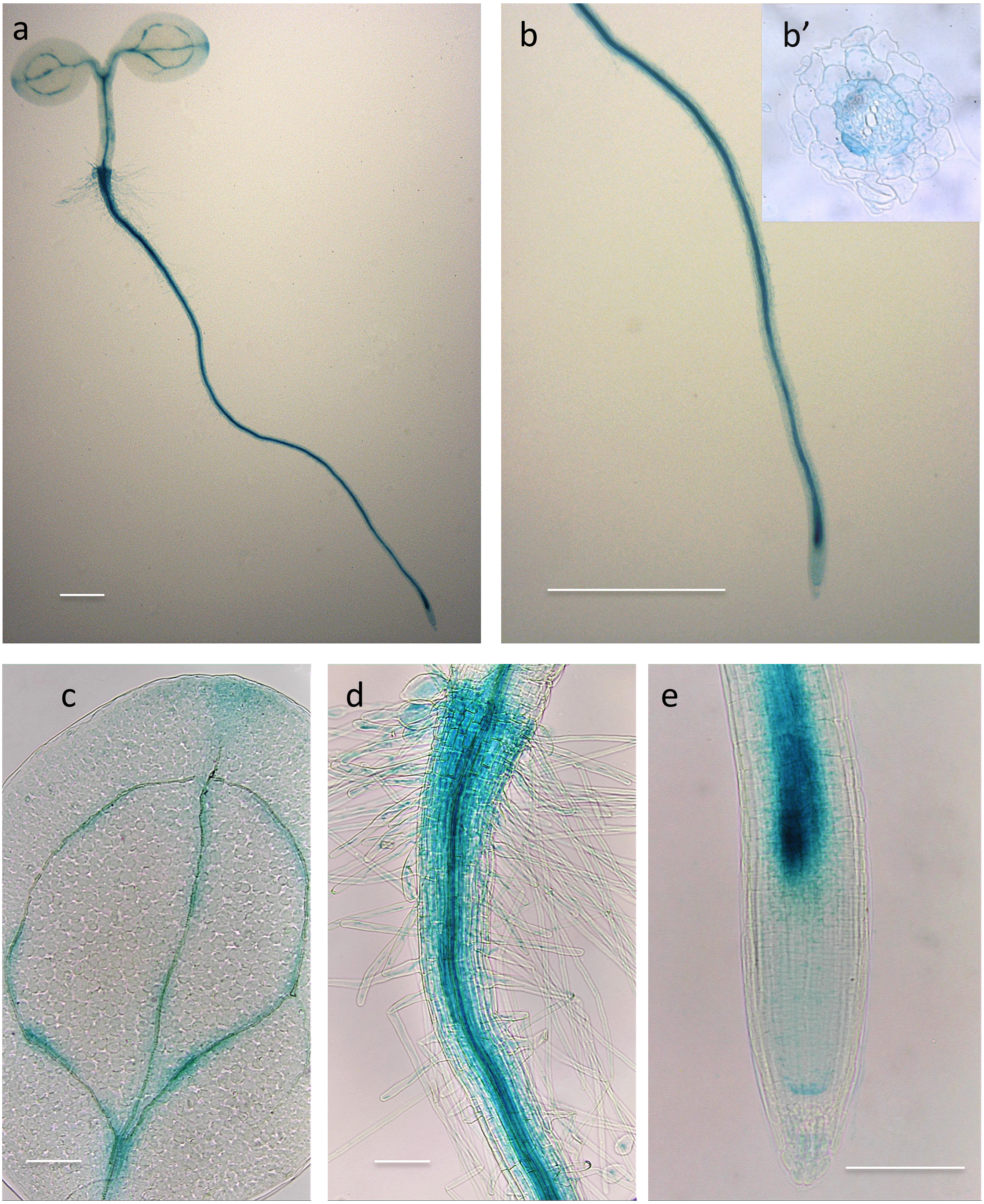

f

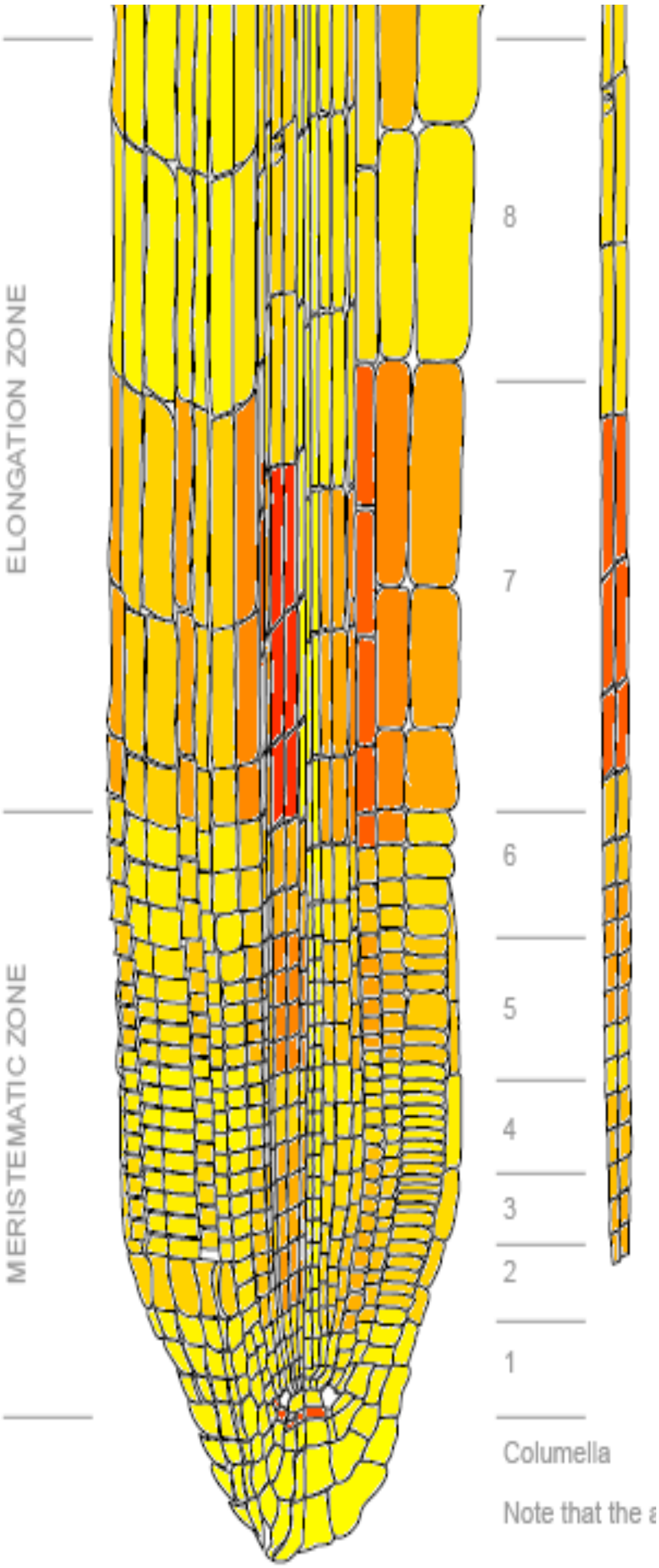


bar.utoronto.ca.

(a-e) *proVAMP714::GUS* is expressed in vascular tissues.

(a) Whole seedling, 4 dpg, bar = 1 mm.

(b) Seedling root, 4 dpg with transverse section in mature region of root, showing GUS activity in the stele (b'), bar = 1 mm.

(c) Cotyledon at 4 dpg, bar = 1 mm.

(d) root-hypocotyl junction at 4 dpg, bar = 1 mm.

(e) primary root tip, 4 dpg, bar = 100 µm.

(f) Expression heat map of VAMP714 gene in primary root of Arabidopsis.

Visualized using online tool at http://bar.utoronto.ca/eplant/. Red denotes high expression, yellow denotes low expression.

**Exocytic and endosomal cycling**

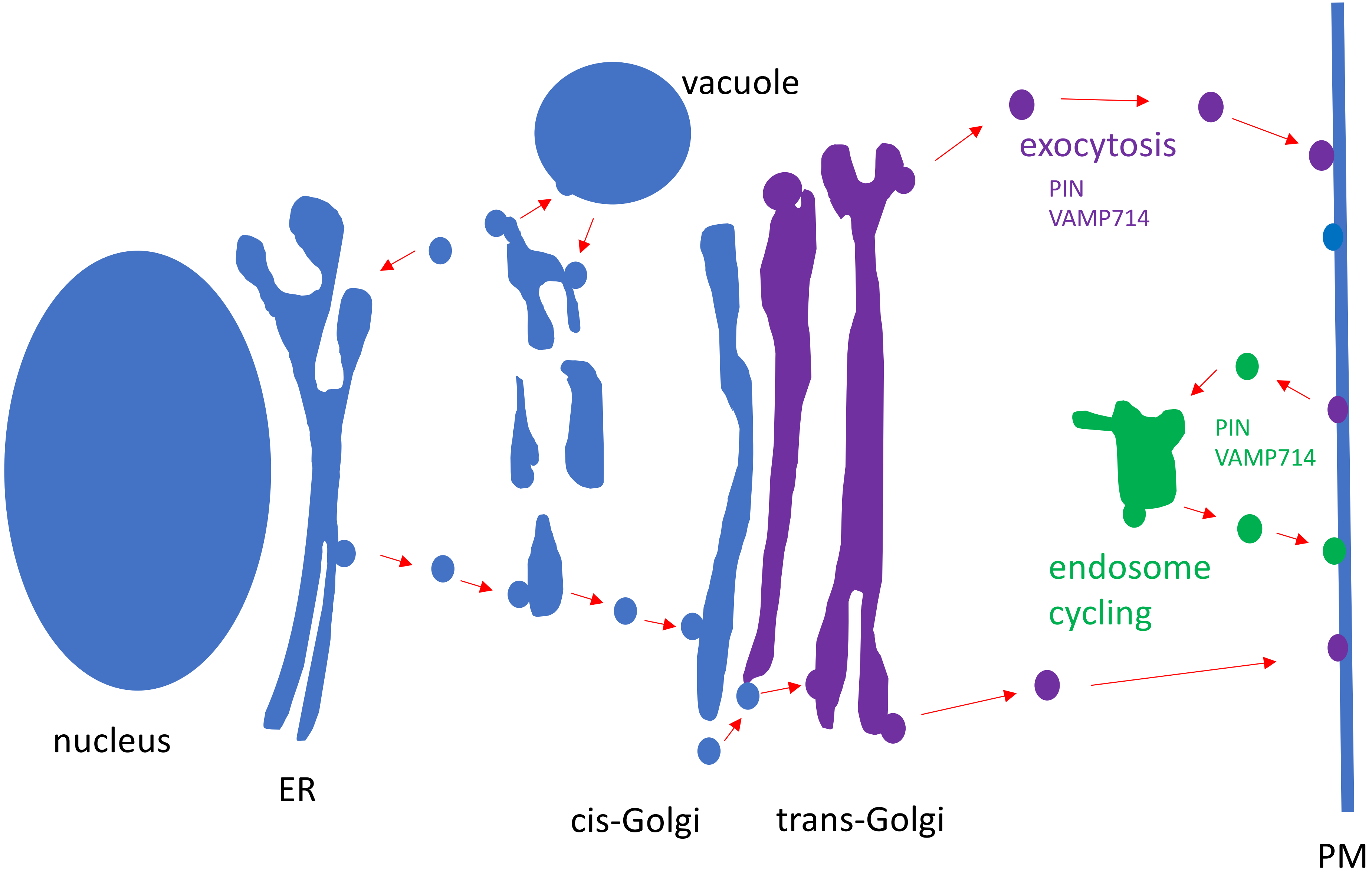


Suppl. Fig. 3

Our data show that VAMP714:mCherry-positive vesicles (purple) move towards the plasma membrane, and co-localize at the plasma membrane (PM) with PIN proteins. VAMP714 also accumulates in BFA bodies and in aggregates following latrunculin B treatment, in the same manner as PIN proteins, both processes being part of endosome recycling (green). VAMP714 is also required for PIN1-positive BFA body formation. It is therefore proposed that VAMP714 is required for both exocytosis of PIN vesicles to the plasma membrane and for PIN cycling between the plasma membrane and endosomes, a process sensitive to BFA and latrunculin B in Arabidopsis.
